# What EEG doesn’t see: a whole-brain MEG map of sleep spindles and slow oscillations

**DOI:** 10.64898/2026.10.09.758064

**Authors:** Anne Calvel, Keelin Greenlaw, Christopher J. Steele, Emily B.J. Coffey

**Affiliations:** Concordia University, 7141 Sherbrooke St W, Montreal, H4B 1R6, Canada; Montreal Neurological Institute, McGill University, 3801 Rue University, Montreal, H3A 2B4, Quebec, Canada

**Author notes:** Corresponding author (A. Calvel); (E.B.J. Coffey).

## Abstract

Sleep spindles and slow oscillations (SOs) are hallmarks of non-rapid eye movement (NREM) sleep, linked to memory consolidation and restorative functions. They are usually detected in one or a few scalp electroencephalography (EEG) channels with parameters that vary across studies, and serve as the reference even in magnetoencephalography (MEG), fMRI and intracranial studies. We argue that these choices filter which neural activity is captured. Nineteen young adults took a daytime nap during simultaneous MEG and EEG. Using MEG source imaging across 416 cortical, subcortical and cerebellar regions, we mapped the amplitude and density of SOs (0.5–4 Hz, and 0.5–2 and 2–4 Hz separately) and of spindles (9–18 Hz, and slow, 9–12.9 Hz, and fast, 13–18 Hz, separately), and related each region’s activity to a single scalp derivation (Cz–M1). Both rhythms were detected throughout the brain, including the thalamus, hippocampus, amygdala, basal ganglia and cerebellum. The balance between bands varied across regions: fast spindles predominated over slow spindles along the dorsal midline, around the central sulcus and in the thalamus, slow spindles predominated in occipital cortex and the cerebellar hemispheres, and 2–4 Hz SOs were relatively most prominent in occipital cortex. Amplitude and density formed reliable but distinct maps: regions with larger fast spindles also had more of them, whereas SOs, and slow spindles once source depth was controlled, were larger where they were less frequent. More SOs were detected in every region than at Cz–M1, whereas more fast spindles were detected at Cz–M1 than in any single region. Around the SO, slow-spindle activity had the same timing at the scalp and in the sources, but the up-state increase in fast-spindle activity seen at the scalp was absent from local sources. Slow spindles were the most strongly shared events, and sensorimotor sources the most scalp-like. Thus, a single scalp derivation offers a selective and partly composite view of sleep oscillations, with implications for interpreting their functional roles and for targeting them with closed-loop auditory stimulation.

**Highlights:**

- Whole-brain MEG source maps of NREM slow oscillations and sleep spindles
- Amplitude and density maps align for fast spindles but diverge for slower events
- More slow oscillations are detected in every region than at the scalp
- More fast spindles are detected at the scalp than in any single region
- Fast spindles in sensorimotor cortex are the most scalp-like
- Scalp up-state fast-spindle activity is absent from local sources

## 1. Introduction

Slow oscillations (SOs), characterized by coordinated transitions of large neuronal populations between up- and down-states, and sleep spindles, brief waxing-and-waning oscillations, are two hallmarks of non-rapid eye movement (NREM) sleep. Both have been associated with memory consolidation (Fernandez and Lüthi, 2020; Kumral et al., 2023) and next-day cognitive performance (Lafortune et al., 2013; Ferrarelli et al., 2019). SOs have further been linked to restorative functions of sleep, including synaptic homeostasis and immune and autonomic regulation (Tononi and Cirelli, 2006; Besedovsky et al., 2017; Grimaldi et al., 2019). Sleep spindles are commonly classified into slow and fast subtypes based on their frequency, with the two subtypes showing distinct predominant topographies. Fast spindles have been associated with declarative memory consolidation (Cairney et al., 2018; Denis et al., 2021) and, in some studies, with procedural memory consolidation (Barakat et al., 2011; Lustenberger et al., 2016). Slow spindles predominate in frontal regions, but their specific contribution to memory processing remains less clearly established (Mölle et al., 2011; Cox et al., 2017). Growing evidence suggests that the precise coupling between SO phase and spindle amplitude plays an important role in declarative memory consolidation (Mikutta et al., 2019; Muehlroth et al., 2019; Hahn et al., 2020).

The active systems consolidation theory of memory provides a framework for understanding how this coupling supports memory consolidation. According to this theory, cortical SOs provide top-down signals that drive fast thalamocortical spindles, creating windows for the gradual transfer of memory-related information from the hippocampus to the neocortex for long-term memory storage (Diekelmann and Born, 2010; Klinzing et al., 2019).

However, evidence from high-density EEG, MEG, fMRI, and intracranial recordings suggests that this model does not capture the full complexity of SOs and spindles across the brain. Even though spindles are conventionally divided into slow and fast classes, their frequency seems to vary more continuously across cortical regions (Gonzalez et al., 2022). Spindle activity may also extend beyond the classical thalamocortical network. Indeed, intracranial recordings have revealed spindle-frequency oscillations in the human basal ganglia (Salih et al., 2009). Cerebellar spindle activity has also been reported through direct recordings in non-human primates (Xu et al., 2021) and, more recently, source-localized MEG in humans (Greenlaw et al., 2026), raising the possibility of a local cerebellar contribution to spindle generation. SOs show similar complexity in both their generation and spatial dynamics. SOs recorded in the anterior thalamus precede their neocortical counterparts by approximately 50 ms, suggesting that the anterior thalamus may contribute to their initiation (Schreiner et al., 2022). Across the cortex, SOs can occur locally or across widespread regions (Fattinger et al., 2025), and can propagate as travelling waves (Massimini et al., 2004; Hangya et al., 2011). This spatial complexity also extends to spindle– SO coupling, which shows a systematic phase precession along the anterior–posterior axis of the human brain (Bender et al., 2025).

Historically, SOs and spindles were identified visually by a trained sleep technician, generally in one or a few EEG electrode channels positioned on the dorsal or frontal scalp as with common polysomnography electrode montages. Amplitude and frequency criteria were shaped by what a human scorer could see easily by eye, and subsequently developed automatic detection algorithms use these visually detected or detectable events as the reference standard (Lacourse et al., 2019). EEG-detected events therefore remain the reference even in studies using fMRI, MEG, or intracranial recordings: events are first identified in EEG, and activity occurring at the same time is then examined in the other modality (Jourde et al., 2024; Klinzing et al., 2016; Jegou et al., 2019).

However, there is little consensus on the algorithm parameters used to detect SOs and spindles. SO detection criteria differ across studies in both frequency and amplitude. For example, Carrier et al. (2011) defined slow waves as activity below 4 Hz with a peak-to-peak amplitude above 75 *µ*V, Massimini et al. (2004) used a 0.1–4 Hz band and a peak-to-peak threshold of 140 *µ*V, Mölle et al. (2009) filtered activity below 2 Hz and used participant-specific amplitude thresholds, and Ladenbauer et al. (2017) used a 0.16–1.25 Hz band and retained the 25% highest-amplitude events. Spindle detection criteria also vary in the literature. For example, Martin et al. (2013) detected 11–15 Hz activity when its RMS envelope exceeded the 95th percentile for at least 0.5 s, Staresina et al. (2015) detected 12–16 Hz activity when its RMS envelope exceeded the 75th percentile for 0.5– 3 s. Slow and fast spindle boundaries also vary, ranging from 9–12 and 12–15 Hz (Mölle et al., 2011) to 8–13 Hz and 13–18 Hz (Bergamo et al., 2025).

Furthermore, interest is high in real-time neuromodulatory techniques such as closed-loop auditory stimulation (CLAS). Due to the need to detect an event before it is complete so as to be able to interact with it, the criteria used for detection must be based on the characteristics of the event’s onset. Consequently, criteria that require the complete event cannot be used, such as spindle duration or SO trough-to-subsequent-peak amplitude when stimulation is delivered during the rising phase. The effectiveness of the onset-based criteria must therefore be carefully validated against offline detection approaches (Valenchon et al., 2022). Another approach circumvents the need to agree on detection parameters by calculating the spectra over a window of time (generally several seconds or a traditional 30 s sleep scoring epoch), and extracting a mean amplitude within a frequency range of interest. Given the large changes in 1/f (fractal) activity over states of consciousness, researchers sometimes separate the spectra into fractal and oscillatory components so as to be able to better observe truly oscillatory activity in SO and spindle bands (Greenlaw et al., 2026). All of the spectral methods confound amplitude and density rather than fully avoiding the problem: e.g., a similar result can be obtained by having fewer high amplitude spindles or more low amplitude spindles within a given unit of time. Results generated using these methods can be difficult to compare with a larger body of EEG research, and are less applicable for targeting specific events in real time neuro-modulation approaches.

Detection choices are not neutral. EEG data from a single channel, or even from a larger number of channels, capture a mixture of the subset of neural sources that happen to project onto the scalp differently at the recording and reference electrodes. Frequency ranges and amplitude criteria interact with channel selection and with the properties of the participant: older adults, for example, have lower-amplitude slow waves that may not reach fixed detection thresholds (Carrier et al., 2011). Minor-seeming variations in detection parameters also matter because different processes may occur in different frequency ranges; for example, 2–4 Hz delta waves and slower SOs appear to play different roles in memory (Benoit et al., 2000; Kim et al., 2019). Detection choices therefore filter what is captured in measures of event amplitude or density, and with them our conclusions about the roles of these events in sleep, about population differences, and about how the brain consolidates memory. For example, the best-supported coupling model, in which SO up-states group spindles that in turn group hippocampal ripples (Staresina et al., 2015; Klinzing et al., 2019), rests largely on events defined at the scalp or in a small number of accessible structures, which leaves open whether it generalizes to other circuits.

There is growing interest in directly modulating sleep oscillations to better understand their functional roles. Among non-invasive approaches, closed-loop auditory stimulation (CLAS) has emerged as a promising technique that uses realtime detection of sleep oscillations to deliver precisely timed auditory stimuli, enabling direct interaction with ongoing sleep oscillations (Ngo et al., 2013). Depending on stimulation timing, CLAS can enhance or disrupt endogenous sleep oscillations (Jourde et al., 2025), providing a means of causally probing their roles and advancing our understanding of sleep’s contributions to health and cognition.

Beyond its value as a research tool, CLAS holds promise for a range of clinical applications. Notably, recent studies have explored its potential to improve memory consolidation in patients with mild cognitive impairment (Papalambros et al., 2019) and Alzheimer’s disease (López-Larraz et al., 2026). CLAS has been shown to reduce interictal epileptiform activity in children with epilepsy (Wong et al., 2026). Beyond these neurological applications, CLAS has also been shown to modulate physiological processes relevant to health, including immune responses, endocrine regulation (Besedovsky et al., 2017) and autonomic function (Grimaldi et al., 2019).

Despite advances in neuroimaging, sparse-montage EEG remains the most widely used approach for monitoring sleep and is likely to remain so due to its accessibility and practicality. However, understanding which neural sources are captured or missed by conventional EEG detection methods, and how activity originating from different brain regions is represented at the scalp, is essential both for interpreting existing findings and for developing more precise and effective targeted interventions.

Magnetoencephalography (MEG) can resolve activity not only across the cortex but also, with appropriate source modelling, in deep and cerebellar structures (Greenlaw et al., 2026; Rattray et al., 2025; Pizzo et al., 2019; López-Madrona et al., 2022), making it possible to detect SOs and spindles at their sources. Here, nineteen young adults took a daytime nap with simultaneous MEG and EEG, and we detected SOs and slow and fast spindles in each of 416 cortical, subcortical and cerebellar regions. We asked three questions. First, how do SO and spindle amplitude and density vary across the brain, including deep structures? Second, are amplitude and density two views of one spatial pattern, or distinct properties? Third, how is each region’s activity represented at the scalp? For the third question we used a single central derivation, Cz–M1, because single central channels are typical of both polysomnography and closedloop stimulation, and compared each region with the scalp in three ways: the correlation of continuous signals, the coincidence of detected events, and the timing of spindle-band activity around SOs. Together, these analyses provide a whole-brain picture of where sleep’s defining oscillations occur and how they appear at the scalp, as a basis for interpreting their functional roles and for targeting them with tools such as closed-loop auditory stimulation.

## 2. Methods

### 2.1 Participants

Twenty neurologically healthy young adults were recruited; one was excluded because a metal dental appliance caused magnetic artefacts, leaving a final sample of 19 (13 females; 18 right-handed; mean age = 21.7 years, SD = 2.8). Note that this is the same data set as described in Greenlaw et al. (2026). The sample size is comparable to or larger than those of previous MEG studies of sleep neurophysiology (Brancaccio et al., 2020; Sjøgård et al., 2025; Weber et al., 2021). Participants had normal or correctedto-normal vision and reported no history of neurological or sleep disorders. All participants provided informed consent and received financial compensation for their participation. The study procedures were approved by the Research Ethics Boards of the Montreal Neurological Institute and Concordia University.

### 2.2 Study design

The data were collected within a larger project on neuroplasticity across states of consciousness. Only the nap recordings are analyzed here; the complete protocol, which also included resting-state and motor learning recordings, is reported in Greenlaw et al. (2026). In brief, to raise sleep pressure, participants went to bed one hour later than usual on the night before the session and abstained from caffeine on the morning of the session. Each session started at about 12:00, placing the nap opportunity in the mid-afternoon dip in alertness (Milner and Cote, 2009). After giving informed consent and changing into sleepwear, participants completed questionnaires on sleep habits and possible sleep disorders (Pittsburgh Sleep Quality Index, PSQI, Buysse et al., 1989; Munich Chronotype Questionnaire, MCTQ, Roenneberg et al., 2003), a Subjective Fatigue Scale, a custom health questionnaire screening for conditions that could affect sleep, and the Montreal Music History Questionnaire (MMHQ; Coffey et al., 2011), which documents musical training. For the recording, participants lay supine in the magnetically shielded room. To maximize coverage of posterior and inferior regions, including the cerebellum (Andersen et al., 2020), the head was inserted as far as possible into the MEG helmet with the chin tilted slightly down, and foam padding was used to limit head and body movement. Resting-state recordings and a motor sequence learning task were followed by a 2.5-hour nap opportunity in darkness, during which participants were asked to rest even if they did not fall asleep. At the end of the nap period, participants were woken gently and given time to become fully alert before subsequent recordings. A T1-weighted anatomical MRI was acquired in a separate session for MEG source reconstruction.

### 2.3. Data acquisition

MEG and polysomnographic signals were recorded simultaneously with a 275-channel whole-head CTF MEG system with integrated EEG inputs (Omega 275, CTF Systems Inc.). Scalp EEG electrodes were placed at Cz, C3 and C4 (10–20 system) and at both mastoids, with the ground on the forehead and the left mastoid (M1) as the online reference. Bipolar electrode pairs recorded chin and neck EMG, EOG (electrodes beside the eyes) and ECG (chest), to support sleep scoring and the identification of ocular and cardiac artefacts in the MEG. All signals were sampled at 2400 Hz. Each recording block was saved as a separate run: 5 min for each resting-state recording, 12 min for each motor-task recording, and five 30-min runs for the nap. Head position was measured before each run from three head-position indicator coils (forehead, left and right), whose locations, together with the head shape, had been digitized with a Polhemus Isotrak system (Polhemus Inc., VT, USA) for co-registration with the anatomical MRI. T1-weighted anatomical images were acquired on a 3 T Siemens scanner with an MPRAGE sequence (176 slices; field of view 256 × 256 mm; TR/TE/TI = 2300/2.98/900 ms; 1 mm isotropic voxels) and used for subject-specific MEG source modelling.

### 2.4. Data pre-processing

Pre-processing was carried out in Brainstorm (Tadel et al., 2011) and with custom MATLAB scripts (The MathWorks Inc., MA, USA).

#### Sleep scoring

For scoring, EEG (C3 and C4, referenced to M1) was band-pass filtered at 0.1–20 Hz, EOG at 0.1– 5 Hz and EMG at 10–58 Hz, and all signals were resampled to 150 Hz. Each 30-s epoch was visually classified as wake, N1, N2, N3 or REM sleep following AASM criteria (Iber et al., 2007). Each epoch was scored by one researcher and independently reviewed by a second. Epochs in which movement or artefacts prevented scoring were not classified as sleep, so the N2 and N3 epochs entering the analyses were free of major artefacts.

#### EEG for MEG–EEG comparisons

Analyses relating MEG to scalp EEG used a single derivation, Cz referenced to the left mastoid (Cz–M1), which was of good quality in all participants. Cz lies on the midline a few centimetres from C3 and C4 and records largely the same activity, so additional central channels would add little independent information. A single central derivation was also chosen because it approximates the approach in closed-loop auditory stimulation applications, which typically detect events in one dorsal or frontal channel (with a mastoid, earlobe, or occipital reference). Cz was band-pass filtered at 0.1–20 Hz and resampled to 150 Hz.

#### MEG

Ocular and cardiac artefacts were attenuated with signal-space projection in Brainstorm (Tesche et al., 1995): projectors explaining at least 12% of the signal were removed if their topography matched ocular or cardiac activity on visual inspection. MEG data were then band-pass filtered at 0.1–37.5 Hz and resampled to 150 Hz.

### 2.5. Source reconstruction and signal extraction

Source reconstruction and signal extraction followed the pipeline described in Greenlaw et al. (2026), and the same source estimates were used here. In brief, each participant’s T1-weighted image was processed with FreeSurfer (Fischl et al., 2002; Fischl, 2012) to reconstruct the cortical surface and segment subcortical structures, including the thalamus, brainstem and cerebellum. The segmentation and cortical surface were imported into Brainstorm and combined into a volume source space with a 5 mm isotropic grid spanning all structures. A head model was computed for each run with the overlapping-spheres method, and sensor noise was characterized from 2-min empty-room recordings acquired before each session. Source activity was estimated with minimum norm estimation (MNE) using unconstrained source orientations and Brainstorm’s default parameters (Tadel et al., 2011; Gramfort et al., 2014), except that depth weighting was increased from 0.5 to 0.7 to improve sensitivity to deep sources (Lin et al., 2006; Gorina-Careta et al., 2021).

The source space was parcellated into 416 ROIs: 340 cortical and 44 subcortical regions (including the hippocampus and amygdala) from the AICHA atlas (Joliot et al., 2015), and 32 cerebellar regions, including the vermis, from AAL3 (Rolls et al., 2020). AICHA was chosen because its parcels are relatively small and functionally homogeneous. The atlases, defined in MNI space, were mapped onto each participant’s anatomy in Brainstorm using an affine registration to the MNI ICBM152 template (Ashburner and Friston, 2005). For each ROI and 30-s epoch, three time series were extracted, one for each source orientation (x, y and z), each the mean across the ROI’s grid points.

Retaining the three orientations, rather than projecting onto a single dominant orientation, avoids assuming a fixed source orientation, which is not well defined for subcortical and cerebellar structures or for ROIs that span folded cortex; the three orientation signals were combined at the detection stage (Section 2.6). Averaging across grid points can partly cancel activity with opposing orientations within an ROI, which is one reason for using an atlas with relatively small parcels; it may reduce amplitude estimates in large or highly folded regions. Despite these limitations, regionspecific MEG signals have been recovered from deep and cerebellar structures with comparable approaches, including the hippocampus and amygdala (Attal and Schwartz, 2013; Dumas et al., 2013), thalamus (Rattray et al., 2025), subcortical auditory nuclei (Coffey et al., 2016) and cerebellum (Samuelsson et al., 2020; Andersen et al., 2020), and in this dataset cerebellar signals were distinguishable from neighbouring occipital cortex (Greenlaw et al., 2026).

### 2.6. Detection methodology

SOs and spindles were detected independently in each ROI and each 30-s epoch of N2 and N3 sleep, using adaptations of established scalp-EEG methods (Mölle et al., 2011; Muehlroth et al., 2019). In contrast to most scalp detectors, which set a single threshold per channel across the whole recording, all thresholds here were defined relative to the signal of the same ROI within the same epoch. This makes detection robust to differences in absolute signal strength between ROIs (e.g., due to source depth), between runs (head position) and over the course of the nap, but it means that density reflects how often activity stands out from the local background rather than its absolute magnitude, and that events are found in most epochs (as would be expected in 30 s N2, N3 epochs as they are defined in EEG by their presence). Consequences of this choice for comparison with scalp-based studies are considered in the Discussion.

#### 2.6.1. Slow oscillations

SOs were detected separately in the broad SO band (0.5– 4 Hz), and in the 0.5–2 Hz and 2–4 Hz bands. For each ROI, each of the three orientation signals was band-pass filtered in the detection band (fourth-order Butterworth, zero-phase), and the three filtered signals were summed. The summed signal was mean-centred, and the first and last 0.5 s of each epoch were excluded to avoid edge effects. Zero-crossings were identified on a version of the signal smoothed with a 50-ms moving average, and candidate events were defined by three consecutive zero-crossings delimiting a negative half-wave followed by a positive half-wave. Candidates were retained if (i) their duration was 0.25–2.0 s (0.5–4 Hz band), 0.5–2.0 s (0.5–2 Hz band) or 0.25–0.5 s (2–4 Hz band) and (ii) their peak-to-peak amplitude exceeded 2.5 times the standard deviation of the detection-band signal of the same ROI and epoch.

#### 2.6.2. Source polarity alignment

With unconstrained source orientations, the summed orientation signal of an ROI reflects the dominant direction of current flow in that region, but its sign depends on how the tissue is oriented relative to the arbitrary x, y and z axes of the source space. This does not affect sign-free measures such as RMS envelopes, and has little effect on SO amplitude and density, but it matters for any analysis that aligns MEG and EEG by phase, such as the SO-locked analyses below, where an inverted ROI would be aligned to its up-state rather than its down-state. We therefore aligned the polarity of each ROI to the scalp: for each participant and ROI, 4-s MEG and simultaneously recorded Cz–M1 EEG segments were extracted around each detected MEG SO trough, with the trough centered in the window. These segments were averaged across SO events within each epoch and then across N2 and N3 epochs. Polarity was determined by comparing the sign of the averaged MEG and Cz–M1 EEG signals at the trough. MEG signals classified as inverted were multiplied by −1, and SO detection was repeated on the corrected MEG signals. After polarity correction and SO redetection, the MEG and Cz–M1 waveforms time-locked to the corrected MEG SO troughs were visually inspected to verify that the troughs were aligned in the same direction. SO-locked sigma RMS profiles were also examined as an additional qualitative check of the corrected alignment. Aligning to Cz–M1 fixes a common sign convention for comparison with the scalp; it does not imply that all regions generate SOs of the same polarity as the scalp signal. The proportion of participants requiring inversion in each ROI is shown in Supplementary Fig. S1.

#### 2.6.3. Sleep spindles

Spindles were detected separately in three bands: all spindles (9–18 Hz), slow spindles (9–12.9 Hz) and fast spindles (13–18 Hz), each with the same procedure. For each ROI, each of the three orientation signals was bandpass filtered in the detection band (fourth-order Butterworth, zero-phase), and its RMS envelope was computed in a 0.2-s moving window. The three envelopes were summed to give a single envelope per ROI Because RMS envelopes are sign-free, spindle detection does not depend on source polarity. Spindles were detected when the envelope exceeded the mean plus 1.5 standard deviations of the envelope within the same ROI and epoch, and events lasting 0.5–3.0 s were retained.

### 2.7. Metrics and aggregation

#### 2.7.1. Slow oscillations

For each ROI and 30-s epoch, SO amplitude was the mean trough-to-peak amplitude of the detected events (the difference between the negative peak and the following positive peak of the summed, polarity-aligned orientation signal, often termed peak-to-peak amplitude), and SO density was the number of detected events per minute.

#### 2.7.2. Sleep spindles

For each ROI and 30-s epoch, spindle amplitude was the mean, across detected spindles, of the peak value of the summed RMS envelope during each spindle, and spindle density was the number of detected spindles per minute.

#### 2.7.3. Aggregation across epochs

For each participant and ROI, epoch values were averaged across all N2 and N3 epochs, with each epoch weighted equally. For density, no minimum number of detected events was required, and all participants contributed to all ROIs. For amplitude, at least one detected event per ROI was required for a participant-level value to be defined.

### 2.8. Group-level maps

For each ROI, the group mean and median across participants were computed. The resulting group maps were then min–max normalized across ROIs to the range [0,1] for visualization. Group maps show the median across participants, as in Greenlaw et al. (2026), to limit the influence of outlying participants. Maps based on the mean showed similar patterns; examples are given in Supplementary Figs. S2 and S3.

#### Fast–slow difference maps

To show where the balance between the two SO bands (2–4 vs 0.5–2 Hz) and between fast and slow spindles differed across the brain, we computed a normalized difference index for amplitude and for density in each ROI, *I* = (*f* − *s*)/(*f* + *s*), where *f* and *s* are the values for the faster and slower band. The index was computed for each participant and the group median taken. The index ranges from −1 to +1, is zero when the two bands are equal, and is independent of the overall level of activity in a region. We used it rather than the raw difference *f* − *s* because raw differences largely reproduced the overall amplitude or density maps (e.g., across ROIs, the raw SO amplitude difference correlated at *ρ* = −0.95 with mean SO amplitude), whereas the normalized index isolates the relative contribution of each band. Because the faster band was consistently smaller or more frequent than the slower band for some measures, each map was centred on its brainwide mean before display, so that the colour scale shows where the balance between bands departs from the brainwide average; brain-wide means are reported in the figure legends. For interpretation, index values were converted to fast/slow ratios, *f* /*s* = (1 + *I*)/(1 − *I*).

### 2.9. Comparison of amplitude and density maps

To test whether amplitude and density capture distinct spatial properties of SO and spindle activity, rather than noisy estimates of a single underlying property, we compared their whole-brain maps separately for each band (SOs at 0.5–4, 0.5–2 and 2–4 Hz; all, fast and slow spindles). Amplitude was taken from the depth-weighted MNE source estimates without normalization to background activity. Because detection thresholds were defined relative to each ROI’s own signal, density does not depend on a region’s absolute signal strength. Group maps were the median across participants. All correlations were Spearman correlations across ROIs, chosen because amplitude and density were skewed across ROIs and because the question concerned the correspondence of regional rankings rather than a linear relationship.

#### Reliability and correction for attenuation

A low correlation between two maps could reflect either a real difference or unreliable measurement. We therefore estimated the reliability of each group map by split-half resampling: participants were randomly divided into two halves (9 and 10), a group map was computed from each half, and the two half-maps were correlated. This was repeated 1,000 times; the correlations were averaged via Fisher’s *z* and corrected to full-sample reliability with the Spearman–Brown formula (Spearman, 1910; Brown, 1910). To estimate the correlation expected in the absence of measurement noise, the amplitude–density correlation was corrected for attenuation by dividing it by the square root of the product of the two reliabilities (Spearman, 1904). A corrected value near ±1 indicates that the maps differ only by noise. The attenuation and Spearman–Brown corrections were derived for Pearson correlations and were applied here as approximations. A 95% confidence interval for the corrected correlation was obtained by bootstrapping participants (1,000 resamples, with both reliabilities re-estimated within each resample from 100 random splits). Because neighbouring ROIs are not statistically independent, correlations across ROIs are reported as descriptive statistics, and inference rests on the participant-level test below.

#### Participant-level test

For each participant, we compared their amplitude and density maps with leave-one-out group maps computed from the remaining 18 participants. Same-measure similarity was the mean Fisher-*z* correlation between the participant’s amplitude map and the group amplitude map, and between their density map and the group density map. Cross-measure similarity was the mean Fisher-*z* correlation between the participant’s amplitude map and the group density map, and vice versa. Same- and cross-measure similarity were compared with paired *t*-tests across participants (*df* = 18), with Holm correction across the six bands. If amplitude and density reflected the same spatial pattern, same- and cross-measure similarity would not differ. We also report the mean within-participant correlation between each participant’s own amplitude and density maps. *Control analyses for source geometry*. Although depth weighting reduces the bias of minimum-norm estimates toward superficial sources, it does not fully equalize amplitude estimates across depth, and deeper sources have lower signal-to-noise ratio. Across the whole brain, ROIs may also differ by tissue compartment in cellular architecture, source orientation and MEG sensitivity. We therefore ran two control analyses. First, every correlation was replaced by a partial Spearman correlation controlling for ROI depth, defined as the distance from each ROI’s centroid (computed from the AICHA and AAL3 atlases in MNI space) to the outer surface of the template brain (convex hull). The ranks of both maps were regressed on the rank of depth and the residuals correlated; this applied to the group correlation, the split-half reliabilities, the bootstrap and the participant-level test. Second, to ensure that the relationships were not driven by differences between cortex and subcortical or cerebellar structures, the analyses were repeated within cortical ROIs only (340 ROIs; hippocampus excluded). Holm correction was applied across bands within each analysis.

### 2.10. Relating MEG sources to scalp EEG

Three complementary analyses related activity in each MEG ROI to the scalp derivation Cz–M1, using the polarity-aligned MEG signals (Section 2.6.2): the correlation between continuous signals, the coincidence of detected events, and SO-locked sigma activity.

#### 2.10.1. Signal correlation

For each participant, ROI and 30-s epoch, we computed the Pearson correlation between Cz–M1 and the MEG signal of the ROI. For SOs, both signals were band-pass filtered at 0.5–4 Hz. For spindles, both signals were band-pass filtered at 9–18 Hz and converted to RMS envelopes with the same procedure as for detection, so that the correlation reflects the co-fluctuation of spindle-band power rather than phase, and is independent of source polarity. Correlations were Fisher *z*-transformed, averaged across epochs within each participant and then across participants, and back-transformed to *r* for display.

### 2.10.2. Event coincidence

SOs and spindles were also detected at Cz–M1, with the same algorithms, bands and epoch-wise thresholds as for the MEG ROIs. For each participant, ROI and band, an MEG event and a Cz event were considered coincident if the MEG and Cz troughs occurred within 200 ms for 0.5–2 Hz SOs, 150 ms for 0.5–4 Hz SOs, and 80 ms for 2–4 Hz SOs. Spindles were considered coincident when their onset-to-offset intervals overlapped. Coincidence was quantified in both directions: the percentage of the ROI’s MEG events matched by at least one Cz event, and the percentage of Cz events matched by at least one event in the ROI. Chance levels were estimated by circularly shifting the MEG event times within each epoch 1,000 times and recomputing both percentages, and excess coincidence was defined as the observed minus the mean chance percentage. Excess coincidence was tested against zero across participants for each ROI (one-sample *t*-test, FDR-corrected across ROIs). Participants with fewer than 20 detected MEG events in an ROI did not contribute to the analysis for that ROI (for spindles, 10–19 participants per ROI; for SOs, all 19).

To separate the two factors that shape this pair of measures, we derived two summary quantities per ROI. The relative event rate,

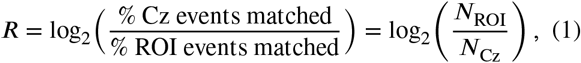

is the ratio of the number of events detected in the ROI to the number detected at Cz, and is positive when the ROI detects more events than Cz. The coincidence strength,

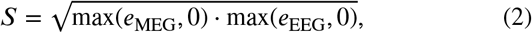

where *e*_MEG_ and *e*_EEG_ are the excess coincidence in the two directions, summarizes how strongly the ROI’s events and the scalp events coincide, independently of which has more. For display, the relative event rate is shown on a logarithmic axis labelled as a ratio.

#### 2.10.3. SO-locked sigma activity

To compare how spindle-band activity is organized around SOs in each region and at the scalp, we computed SO-locked averages of sigma power, rather than counting SO– spindle coupled events. This approach does not depend on spindle detection thresholds or on a coupling window, both of which strongly affected coupling estimates in preliminary analyses, and it can be applied identically to MEG and EEG at the same events.

For each participant and ROI, windows of ±2 s were extracted around each SO trough detected in that ROI (0.5– 2 Hz band), in N2 and N3 sleep. In each epoch, sigma-band RMS envelopes were computed for all (9–18 Hz), slow (9– 12.9 Hz) and fast (13–18 Hz) sigma, with the same filters and RMS window as for spindle detection, both for the ROI’s MEG signal (envelopes computed per orientation and then summed) and for Cz–M1 at the same time points. The SO waveforms themselves were averaged in the same way. For each participant and ROI, envelopes were first averaged across events within each 30-s epoch and then across N2 and N3 epochs. For each ROI, participants with at least one retained SO were included.

Trough and up-state windows were derived from the data. The up-state peak was identified on the group-average MEG SO waveform (each ROI’s waveform scaled to unit trough depth, then averaged across ROIs), and both windows were given a half-width of one quarter of the trough-to-peak interval. For 0.5–2 Hz SOs, the up-state peak occurred 0.40 s after the trough, giving a trough window of −0.10 to 0.10 s and an up-state window of 0.30 to 0.50 s. For each ROI, the up-minus-trough difference in sigma power was tested against zero across participants (one-sample *t*-test, Cohen’s*d*, FDR-corrected across ROIs), separately for MEG and EEG.

For the main figure, normalized curves were averaged within each participant across the ROIs of each region group: sensorimotor cortex (central, precentral and postcentral sulci and paracentral lobule; 34 ROIs), occipital cortex (calcarine, cuneus, lingual, and inferior, lateral, middle, superior and polar occipital gyri; 50 ROIs), remaining cortex (256 ROIs), thalamus (18), hippocampus and amygdala (6), other subcortical nuclei (caudate, putamen and pallidum; 20) and cerebellum (32). The ROIs in each group are listed in Supplementary Table S1.

MEG and EEG were compared within participants with paired *t*-tests on the mean fast-sigma change in three windows (pre-trough, −0.60 to −0.25 s; trough; up-state) and on the interaction (up-state minus trough, EEG vs MEG). Point-wise differences across time were tested with paired*t*-tests at each sample, FDR-corrected across samples within ±1.2 s.

Latencies were compared between MEG and EEG with the jackknife procedure (Miller et al., 1998; Ulrich and Miller, 2001), which estimates latency on leave-oneparticipant-out averages and is more reliable than per-participant latencies on noisy curves. The onset of the pre-trough fast-sigma rise was defined as the time at which the curve first reached half of the rise from its minimum (between −1.0 and −0.4 s) to its maximum (between −0.4 and 0.1 s); the slow-sigma peak was searched between −0.5 and 0.2 s and the SO trough between −0.2 and 0.2 s. The latency difference (EEG minus MEG) measured on the grand average was tested with a jackknife-corrected standard error and a *t*-test with *n* − 1 degrees of freedom.

### 2.11. Statistical analysis

All tests were two-sided with *α* = 0.05. Correlations were averaged via Fisher’s *z* transform. Corrections for multiple comparisons used the Benjamini–Hochberg false discovery rate (Benjamini and Hochberg, 1995) across ROIs for whole-brain maps and across time points for time-resolved comparisons, and the Holm procedure (Holm, 1979) across frequency bands where a small family of tests was compared. Because neighbouring ROIs are not statistically independent, statistics computed across ROIs are reported descriptively, and inference is based on tests across participants. Analyses were performed in MATLAB (The MathWorks Inc.) with custom scripts.

## 3. Results

### 3.1 Sleep architecture

Sleep architecture for this cohort has been described previously (Greenlaw et al., 2026). Across participants, N2 duration ranged from 8 to 96 min and N3 duration from 0 to 73 min.

### 3.2. Slow oscillation amplitude and density vary across the brain, including deep structures

SO amplitude varied about 13-fold across ROIs in the 0.5–2 Hz band and 12-fold in the 2–4 Hz band (Fig. 2). Relative to the cortical median, it was largest in the thalamus (1.2–1.4 times) and, within cortex, in the temporal poles, supramarginal and superior parietal cortex and around the central sulcus. It was smallest in the cerebellar hemispheres (about 0.6 times the cortical median). SO density followed a different gradient and varied much less across ROIs (0.5– 2 Hz: 21–26 events/min; 2–4 Hz: 57–74 events/min): it was highest in occipital cortex (calcarine, lingual, occipital pole) and the cerebellar vermis, and lowest in superior, medial and orbital frontal cortex.

**Figure 1:**
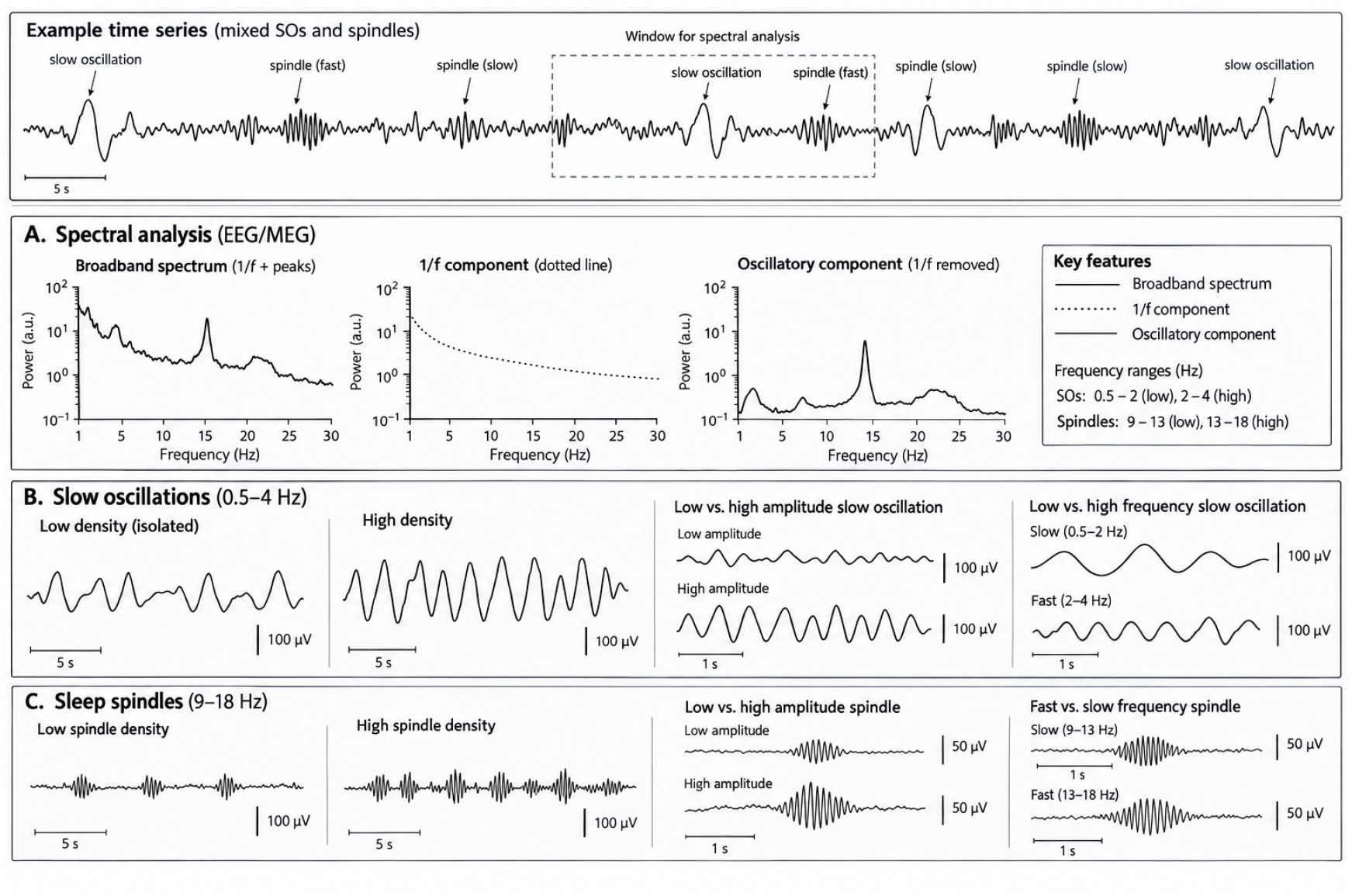
Quantification of NREM sleep microarchitecture. NREM sleep timeseries data (both EEG and MEG) may include slow oscillations and sleep spindles that vary in amplitude and frequency. Their presence can be quantified via spectral analyses (A), in which a window of data is decomposed into power as a function of frequency (the aperiodic component (1/f) is sometimes removed to measure the purely oscillatory component). However, spectral measures mix amplitude and density as data containing few high amplitude events can have a similar spectrum to one containing many low amplitude events. Amplitude and density (illustrated in B and C) can show different relationships to sleep-related processes. Event detection methods can disentangle amplitude and density, but the neural circuits whose activity they each capture may depend on the approach and its parameters, notably filter frequency ranges.

**Figure 2:**
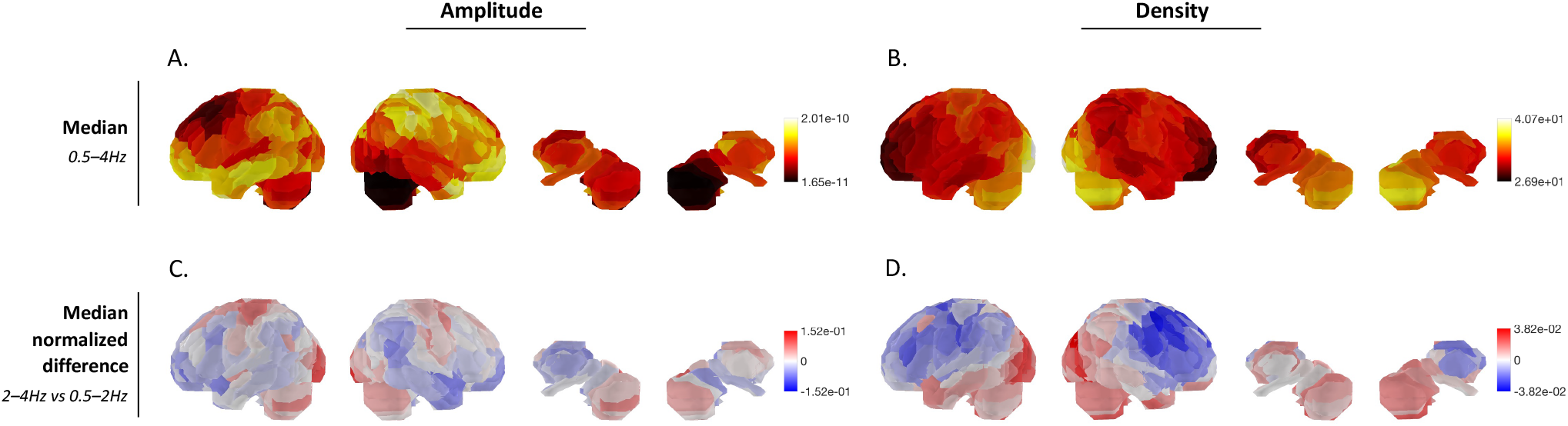
Whole-brain maps of slow oscillation (SO) amplitude and density. (group median across participants, *N* = 19). A: 0.5–4 Hz amplitude, defined as the mean trough-to-peak amplitude of detected SOs. B: 0.5–4 Hz density, defined as the number of detected SOs per minute of N2 and N3 sleep. For A–B, values were normalized within each map; colour bars show the original minimum and maximum values. C: Relative amplitude difference between the 2–4 Hz and 0.5–2 Hz bands. D: Relative density difference between the 2–4 Hz and 0.5–2 Hz bands. For C–D, relative differences were calculated as (*f* − *s*)/(*f* + *s*), where *f* and *s* are the 2–4 Hz and 0.5–2 Hz values. Each map was centred on its brain-wide mean (red: 2–4 Hz relatively more prominent than the brain-wide average; blue: 0.5–2 Hz relatively more prominent). Brain-wide means: amplitude −0.34 (2–4 Hz SOs about half the amplitude), density 0.47 (2–4 Hz SOs about 2.8 times as frequent). Colour scales are set separately for each map; the regional variation in the density ratio is small (fast/slow ratio 2.6–3.1) compared with amplitude (0.42–0.69).

The two SO bands also differed in their spatial distributions (Fig. 2, panels C–D). Across the brain, 2–4 Hz SOs were about 2.8 times as frequent as 0.5–2 Hz SOs, and this ratio varied only modestly across regions (2.6 to 3.1). It was highest in occipital cortex (cuneus, calcarine, lateral and superior occipital) and lowest over frontal and central regions (precentral and central sulci, superior, middle and inferior frontal cortex, anterior cingulate). Amplitude varied more: 2–4 Hz SOs were on average about half the amplitude of 0.5–2 Hz SOs (ratio 0.50), ranging from 0.42 to 0.69 across regions. The two bands were most similar in amplitude in occipital cortex (calcarine, lingual, occipital pole) and the vermis, and most different in the thalamus, orbitofrontal cortex, superior temporal sulcus and anterior insula.

SO densities were higher than values typically reported from scalp recordings, as expected from detection thresholds defined relative to each ROI and epoch (see Discussion).

### 3.3. Spindle amplitude and density

Spindle density showed the expected fast/slow topography (Fig. 3). Fast spindles were most frequent along the dorsal midline (paracentral lobule, cingulate sulcus, mid and posterior cingulate, precuneus) and around the central sulcus (centro-parietal median 1.96 vs frontal 1.46 events/min), and least frequent in occipital cortex and the cerebellum. Slow spindles were most frequent in superior, medial, middle and orbital frontal cortex and the anterior insula (frontal 1.72 vs centro-parietal 1.44 events/min), and least frequent in occipital cortex and the vermis. Spindles of both types were detected in deep structures: densities in the thalamus, hippocampus and amygdala, and other subcortical nuclei were 0.8–1.1 times the cortical median, and lower in the cerebellum (0.56 and 0.70 times for fast and slow spindles).

**Figure 3:**
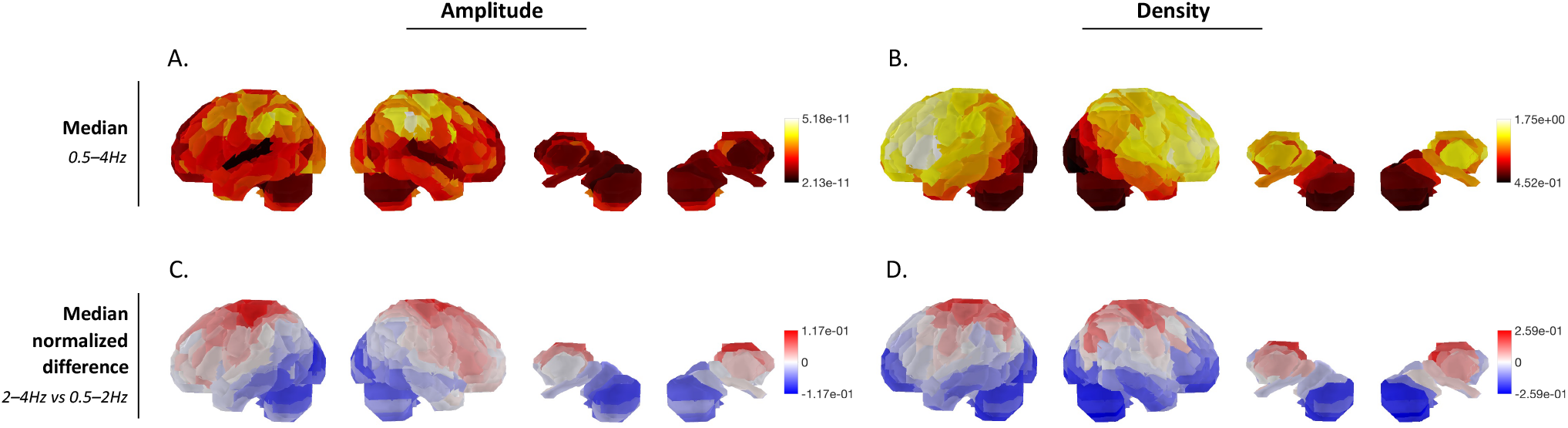
Whole-brain maps of spindles amplitude and density. (group median across participants, *N* = 19). A: 9–18 Hz spindles amplitude, defined as the mean peak RMS amplitude of detected spindles. B: 9–18 Hz density, defined as the number of detected spindles per minute of N2 and N3 sleep. For A–B, values were normalized within each map; colour bars show the original minimum and maximum values. C: Relative amplitude difference between the 13–18 Hz and 9–12.9 Hz bands. D: Relative density difference between the 13–18 Hz and 9–12.9 Hz bands. For C–D, relative differences were calculated as (*f* − *s*)/(*f* + *s*), where *f* and *s* are the fast (13–18 Hz) and slow (9–12.9 Hz) values. Each map was centred on its brain-wide mean (red: 13–18 Hz relatively more prominent than the brain-wide average; blue: 9–12.9 Hz relatively more prominent). Brain-wide means: amplitude −0.12 (fast/slow ratio 0.79; regional range 0.63–0.99), density 0.00 (fast/slow ratio 1.0; regional range 0.6–1.6). Colour scales are set separately for each map.

The balance between fast and slow spindles varied considerably across the brain (Fig. 3, panels C–D). Brain-wide, fast and slow spindles were about equally frequent (ratio 1.0), but the fast/slow density ratio ranged from 0.6 to 1.6 across regions. Fast spindles predominated along the dorsal midline (mid and posterior cingulate, cingulate sulcus, paracentral lobule, precuneus), around the central sulcus and in the thalamus and caudate; slow spindles predominated in occipital cortex, the cerebellar hemispheres and superior frontal cortex.

Spindle amplitude followed a different pattern from density in both bands. It varied about 2.6-fold across ROIs and was largest in lateral parietal cortex (supramarginal, angular, superior and inferior parietal), the thalamus and around the central sulcus, and smallest in anterior cingulate, medial and rostral frontal cortex, superior temporal cortex and cerebellar lobule VI. Fast spindles were smaller than slow spindles almost everywhere (brain-wide ratio 0.79; range 0.63 to 1.0), with the smallest amplitude difference around the central sulcus, paracentral lobule and superior frontal sulcus, and the largest in occipital cortex (cuneus, calcarine, lingual, lateral occipital, fusiform) and cerebellar crus I. Where spindles were largest (lateral parietal cortex) therefore differed from where they were most frequent (the dorsal midline for fast spindles, frontal cortex for slow spindles).

### 3.4 Amplitude and density are distinct properties

Group maps of both measures were highly reliable (splithalf reliability: amplitude 0.91–0.95, density 0.75–0.95; Table 1). Amplitude and density were nevertheless only weakly to moderately related across ROIs, even after correction for attenuation (|*ρ*| ≤ 0.43), so each measure shares at most about 18% of its spatial variance with the other. The direction of the relationship depended on the band. SOs were larger where they were less frequent (0.5–4 Hz: *ρ* = −0.43, 95% CI −0.51 to −0.24; 0.5–2 Hz: *ρ* = −0.41, −0.51 to −0.22), a relationship that was weaker for 2–4 Hz SOs (*ρ* = −0.15,−0.24 to −0.05). In contrast, regions with larger fast spindles also had more of them (*ρ* = 0.35, 0.11 to 0.48). Slow spindles and the combined spindle band showed no reliable relationship in the full ROI set

At the participant level, each person’s amplitude map resembled the other participants’ amplitude maps far more than their density maps, and vice versa. Same-measure similarity ranged from 0.63 to 0.79 and cross-measure similarity from −0.26 to 0.25 (all *t*(18) ≥ 16.1, Holm-corrected *p <* 10^−11^). Within individuals, the amplitude–density correlation followed the group pattern: negative for SOs (0.5–4 and 0.5–2 Hz: mean *ρ* = −0.37), positive for fast spindles (*ρ* = 0.31) and near zero for slow spindles (*ρ* = 0.01).

**Table 1.** Correspondence between amplitude and density maps (all ROIs; *N* = 19 participants, 416 ROIs).

| Band | Rel. amp. | Rel. dens. | $\rho$ (group) | $\rho_{\text{corr}}$ [95% CI] | Same | Cross | $t(18)$ | Within |
| --- | --- | --- | --- | --- | --- | --- | --- | --- |
| SO 0.5–2 Hz | 0.93 | 0.75 | −0.34 | −0.41 [−0.51, −0.22] | 0.63 | −0.22 | 17.4 | −0.37 |
| SO 2–4 Hz | 0.95 | 0.95 | −0.14 | −0.15 [−0.24, −0.05] | 0.79 | −0.11 | 17.2 | −0.12 |
| SO 0.5–4 Hz | 0.94 | 0.83 | −0.38 | −0.43 [−0.51, −0.24] | 0.69 | −0.26 | 21.5 | −0.37 |
| Spindles, slow (9–12.9 Hz) | 0.91 | 0.91 | −0.20 | −0.22 [−0.39, 0.08] | 0.68 | −0.15 | 16.1 | 0.01 |
| Spindles, fast (13–18 Hz) | 0.94 | 0.95 | 0.33 | 0.35 [0.11, 0.48] | 0.76 | 0.25 | 18.0 | 0.31 |
| Spindles, all (9–18 Hz) | 0.92 | 0.89 | −0.04 | −0.04 [−0.25, 0.23] | 0.71 | −0.04 | 21.2 | 0.09 |

Rel. amp., Rel. dens.: split-half reliability of the group (median) map, Spearman–Brown corrected. $\rho$ (group): Spearman correlation across ROIs between group amplitude and density maps (descriptive; ROIs are not independent). $\rho_{\text{corr}}$ : corrected for attenuation; 95% CI from 1,000 participant bootstraps. Same, Cross: mean similarity of each participant's maps to leave-one-out group maps of the same or the other measure (back-transformed from Fisher $z$ ). $t(18)$ : paired test of same vs cross; all Holm-corrected $p < 10^{-11}$ . Within: mean correlation between each participant's own amplitude and density maps.

These relationships were not explained by source geometry. As expected given the lower MEG sensitivity to deep sources, absolute amplitude decreased with source depth, more for spindles (*ρ* = −0.46 to −0.52) than for SOs (*ρ* = −0.24 to −0.28), whereas density was less related to depth (SOs *ρ* = 0.23 to 0.27; spindles *ρ* = −0.29 to 0.06). Controlling for depth did not remove the amplitude–density relationships (Supplementary Table S2). After partialling depth, the negative relation for slower SOs was preserved (0.5–4 Hz: *ρ* = −0.39, −0.48 to −0.22; 0.5–2 Hz: *ρ* =−0.36, −0.47 to −0.17) and the positive relation for fast spindles strengthened (*ρ* = 0.44, 0.16 to 0.60). Slow spindles now showed a negative relationship (*ρ* = −0.44, −0.60 to −0.08), mirroring fast spindles; in the full ROI set this had been partly masked because both amplitude and density decrease with depth. The weak relation for 2–4 Hz SOs was no longer distinguishable from zero after partialling depth (*ρ* = −0.08, −0.20 to 0.01) or within cortex (*ρ* = −0.05,−0.15 to 0.06). Restricting the analysis to cortical ROIs otherwise gave a similar pattern (SO 0.5–4 Hz *ρ* = −0.36; SO 0.5–2 Hz *ρ* = −0.34; fast spindles *ρ* = 0.37). The participant-level test remained highly significant in both control analyses (all *t*(18) ≥ 15.7, Holm-corrected *p <* 10^−11^).

### 3.5. What the scalp shares with each region

Events detected in each ROI coincided with events detected at Cz–M1 more often than expected by chance in every ROI and band, in both directions (all 416 ROIs, FDR-corrected; Fig. 4). Coincidence was nevertheless partial. For spindles, about one fifth of the events in a typical ROI were matched by a Cz event (median across ROIs: slow, 19.0%; fast, 20.0%; chance 3.0% and 5.3%). For SOs, raw percentages were similar (0.5–2 Hz: 19.3%), but because SOs are dense, chance coincidence was also high (11.3%), and excess coincidence was correspondingly modest.

**Figure 4:**
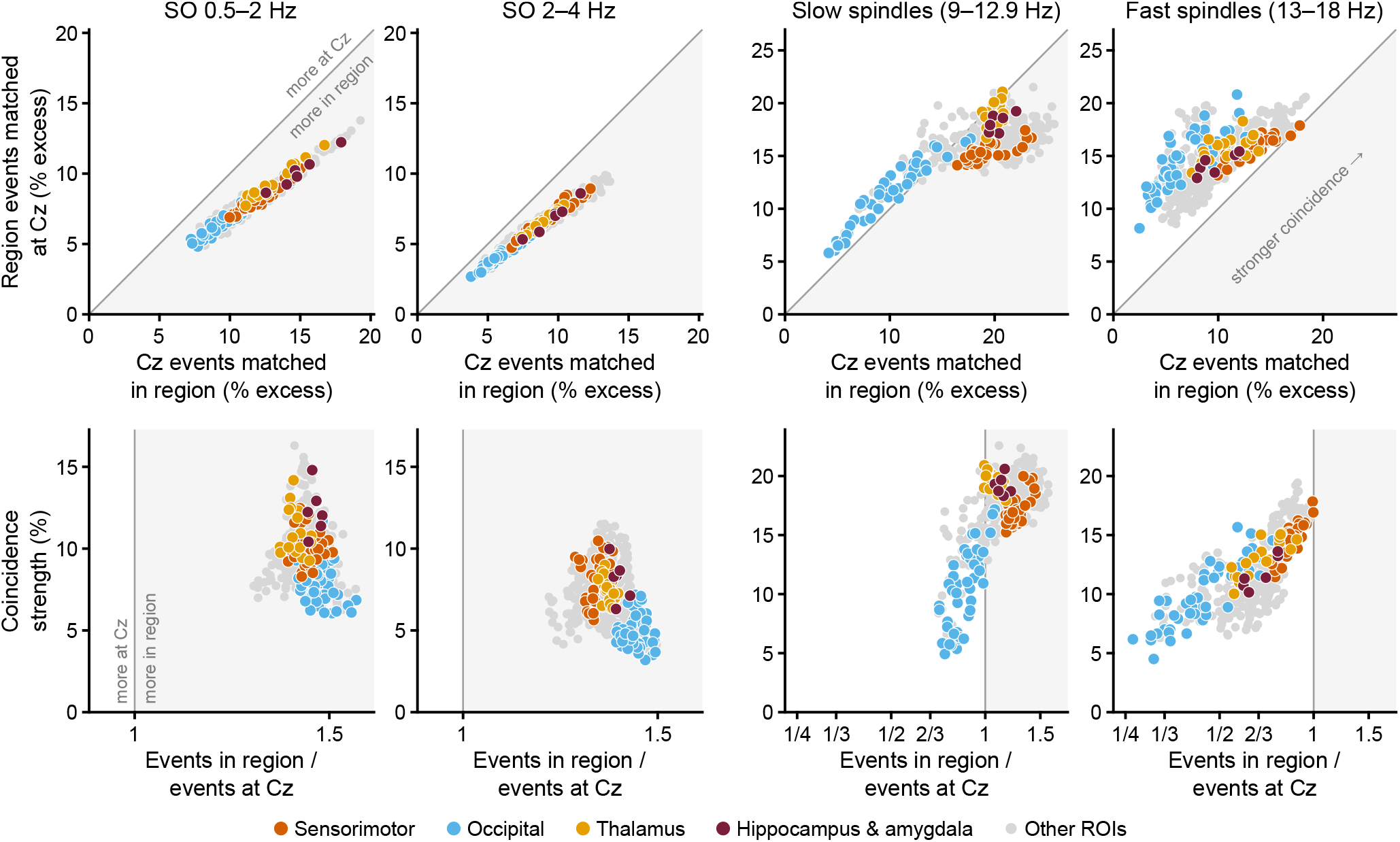
Coincidence of events detected in each region (MEG source) and at Cz–M1, for SOs and slow and fast spindles. More SOs were detected in every region than at the scalp, whereas more fast spindles were detected at the scalp than in any single region; slow spindles were the most strongly shared events. Fast-spindle coincidence was strongest where a region’s event rate approached that of Cz, as in sensorimotor cortex. Each point is one ROI (group mean, *N* = 19); colours as in Fig. 5, other ROIs in grey. Top row: percentage of Cz events matched in the region (*x*) and of the region’s events matched at Cz (*y*), each minus chance (1,000 circular shifts); grey line, identity. All ROIs exceeded chance in both directions (FDR-corrected). Bottom row: relative event rate (events in region / events at Cz) and coincidence strength (geometric mean of the two excess values). Shading marks the side on which more events were detected in the region than at Cz.

Two factors shape these measures: how many events a region produces relative to Cz, and how strongly its events coincide with those at the scalp. SOs and spindles differed in both. In both SO bands, every ROI detected more SOs than Cz, by a similar factor across the brain (0.5–2 Hz: median ratio 1.43, range 1.28 to 1.59; 2–4 Hz: 1.35, 1.20 to 1.50), and regions differed mainly in coincidence strength (0.5–2 Hz: median 9.6%, range 6.1 to 16.3%; 2–4 Hz: 7.0%, 3.2 to 11.5%). Coincidence strength was highest in the hippocampus and amygdala (median 12.1% for 0.5–2 Hz SOs), followed by the thalamus and other subcortical nuclei (10.5 and 10.4%), and lowest in the cerebellum (8.3%), which also detected the most SOs relative to Cz (ratio 1.51). Across ROIs, relative event rate and coincidence strength were weakly negatively related (*ρ* = −0.24 and −0.26 for the two bands).

Spindles showed a different structure, with a sharp contrast between slow and fast spindles. For slow spindles, most ROIs (76%) detected more events than Cz (median ratio 1.17, range 0.71 to 1.59), coincidence strength was high (median 17.8%), and strength increased with event rate before levelling off (*ρ* = 0.63); the thalamus and other subcortical nuclei were among the most strongly coincident regions (19.3 and 19.9%). The cerebellum was the exception: it detected fewer slow spindles than Cz (ratio 0.85) and coincided less with the scalp (14.9%). For fast spindles, no ROI detected as many events as Cz (median ratio 0.68, range 0.26 to 1.00; cerebellum 0.42), coincidence strength was lower (median 11.7%, range 4.5 to 19.4%), and strength rose steeply with event rate (*ρ* = 0.80): the ROIs that detected nearly as many fast spindles as Cz were those whose events coincided most with it.

Continuous signals told a similar story (Supplementary Fig. S4). SO-band activity at Cz–M1 correlated weakly with every ROI (epoch-wise *r* = 0.02 to 0.25), most strongly with frontoventral cortex and the temporal poles, whereas spindle-band power at Cz correlated more strongly and more uniformly with source power (r = 0.19 to 0.51), least in occipital cortex.

### 3.6. SO-locked sigma activity: where MEG and EEG agree and where they differ

At the scalp and in the sources, slow and fast sigma were organized differently around the SO (Fig. 5). Fig. 5 shows four of the seven region groups; all seven are shown in Supplementary Fig. S5, and the statistics below refer to all seven. Slow sigma peaked just before the SO trough and declined into the up-state in every region group, in both MEG and EEG (up-state minus trough, all *p <* 0.03). Its timing agreed closely between signals: slow-sigma peaks in EEG and MEG were within 30 ms of each other in five of the seven groups (47 ms in the thalamus and 80 ms in the hippocampus and amygdala) and differed significantly only for the remaining cortex (27 ms, *p* = 0.028; jackknife; Supplementary Table S3). The amplitude of slow-sigma modulation was also similar: around the trough, EEG and MEG did not differ in five of the seven groups, and EEG was modestly higher in occipital cortex and the thalamus (1.2 and 0.9 percentage points; *p* = 0.002 and 0.025).

**Figure 5:**
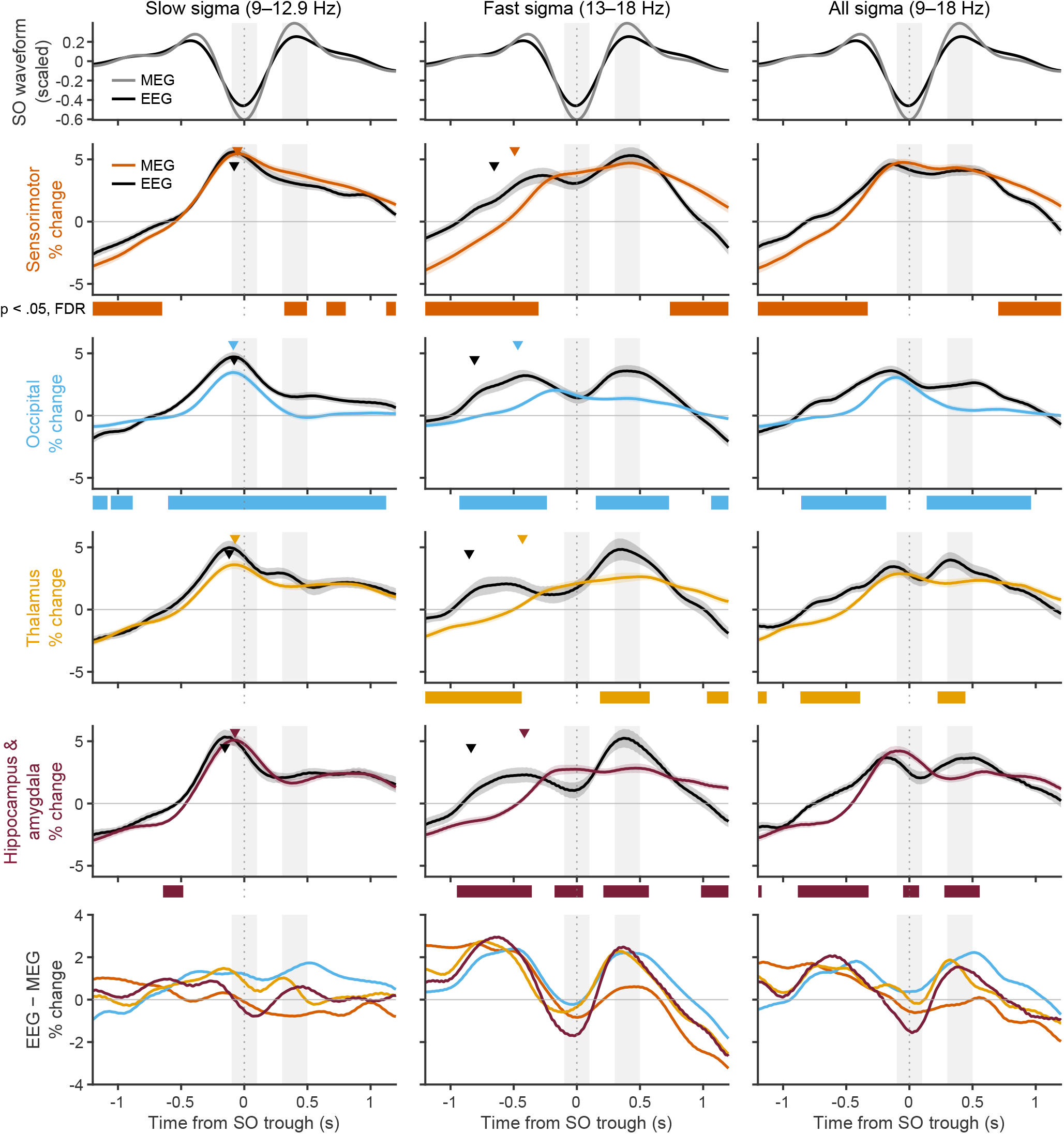
Sigma activity time-locked to SOs (0.5–2 Hz) detected in each region, in the local MEG source and at Cz–M1 at the same events. Slow sigma peaked just before the SO trough in both signals, whereas the up-state rise in fast sigma appeared only at the scalp and began earlier there. Sensorimotor cortex was the most similar to the scalp. Columns: slow (9–12.9 Hz), fast (13–18 Hz) and all (9–18 Hz) sigma. Top row: SO waveform averaged over all ROIs (MEG, grey; EEG, black; each scaled to its range). Rows 2–5: sigma envelope (percent change from its mean over ±2 s) for four region groups; coloured, MEG; black, EEG; shading, ±1 SEM (*N* = 19). Grey bands, trough and up-state windows. Triangles, jackknife latencies (slow sigma: peak; fast sigma: half-maximum of the pre-trough rise). Strips, time points where EEG and MEG differ (FDR-corrected). Bottom row: EEG minus MEG. All seven region groups in Supplementary Fig. S5.

Fast sigma differed systematically between the scalp and the sources. In EEG, fast sigma rose from the trough into the up-state at every region group’s events (up-state minus trough, 1.7 to 3.8 percentage points, all *p* ≤ 0.015), whereas in the local MEG source it did not (0.1 to 0.7 percentage points, all *p >* 0.05); this difference between signals was significant in six of seven groups (interaction, *p* = 0.0002 to 0.048), the exception being sensorimotor cortex (*p* = 0.071). EEG fast sigma also began to rise earlier than MEG fast sigma. The local rise began at a similar time in every group (half-maximum −0.41 to −0.49 s relative to the local trough), whereas the EEG rise at the same events began earlier and was more variable (−0.60 to −0.85 s; Supplementary Table S3). The difference was significant in all groups except the cerebellum (Supplementary Fig. S5), and was smallest in sensorimotor cortex (−0.16 s, *p* = 0.025) and largest in the thalamus and the hippocampus and amygdala (−0.42 s, both *p <* 0.001; jackknife). These offsets were not explained by a shift of the SO itself: at the same events, the EEG SO trough preceded the local trough by only 0 to 27 ms.

The EEG–MEG difference therefore took a consistent form for fast sigma but not for slow sigma (Fig. 5, bottom row). For fast sigma, the difference curves had the same shape in the four groups shown and in the remaining three (Supplementary Fig. S5), with EEG exceeding MEG before the trough and in the up-state and falling slightly below it at the trough; for slow sigma, the differences were small and varied in sign across groups. Finally, combined 9–18 Hz sigma showed no trough-to-up-state modulation in EEG for any region group (all *p >* 0.07), although the local MEG source decreased from trough to up-state in occipital cortex, the remaining cortex and the hippocampus and amygdala (all *p <* 0.001).

## 4. Discussion

We mapped slow oscillations and sleep spindles across 416 cortical, subcortical and cerebellar regions with wholebrain MEG source imaging, and compared each region’s events with those recorded simultaneously at the scalp in a single EEG derivation (Cz–M1). Both rhythms were detected throughout the brain, including deep structures, and their amplitude and density varied across regions. Three further findings stand out. First, with the same detection procedure in both signals, and thresholds set relative to each signal’s own amplitude distribution so that differences in overall signal strength did not determine detection, scalp and source measures disagreed about how often events occur, in opposite directions for the two rhythms: more SOs were detected in every region than at Cz–M1, whereas more fast spindles were detected at Cz–M1 than in any single region. Second, the rise in fast-spindle activity into the SO up-state, regularly observed in EEG recordings, was absent from the local MEG source signal at the same events in every region group, whereas the timing of slow-spindle activity relative to the SO was shared between scalp and sources. Third, the regions where events were largest were not always those where they were most frequent: amplitude and density maps aligned for fast spindles but diverged for SOs and slow spindles, and these relationships held after accounting for source depth. Across analyses, sensorimotor cortex resembled the scalp most closely, and the thalamus and the hippocampus and amygdala differed most in timing. Together, these results suggest that a single scalp derivation offers a selective and partly composite view of sleep oscillations. Below, we consider what the scalp captures, what this implies for SO– spindle coupling, what amplitude and density each reflect, the contribution of deep structures, and the implications for closed-loop stimulation, before turning to limitations and future directions.

### 4.1. What a single scalp derivation captures

Our results indicate that scalp EEG captures only part of the oscillatory activity occurring across the brain (Fig. 4). More SOs were detected in every MEG region than at Cz, by a similar factor across the brain, suggesting that the scalp preferentially records the larger and more widespread SOs and misses many local ones (Nir et al., 2011; Fattinger et al., 2025). Fast spindles showed the opposite pattern: more were detected at Cz than in any individual region, and each region shared only a fraction of them, suggesting that scalp fast spindles combine events from several sources rather than mirroring any single one. Slow spindles behaved as broad, shared events: most regions, including the thalamus, detected at least as many slow spindles as Cz, and their coincidence with the scalp was high.

The correspondence between regions and the scalp formed a gradient rather than a division between “visible” and “invisible” regions. Sensorimotor cortex, close to Cz and rich in fast spindles, was the most scalp-like. Even in this favourable configuration, however, the timing of fast-sigma activity at Cz differed from that in the local sources. Because Cz lies near the centro-parietal maximum of fast spindles, it is arguably the most favourable single derivation for fast spindles; if Cz does not reflect local fast-spindle timing, other derivations are unlikely to do better, although this prediction remains to be tested with full-cap recordings. More broadly, detection parameters may determine which circuits are captured. Distinguishing slower SOs from 2– 4 Hz delta activity (Kim et al., 2019), or slow from fast spindles, may isolate activity from different circuits rather than provide different measurements of a single process. This raises the question of how far the hierarchy of SOs, spindles and hippocampal ripples, established largely from scalp-detected events (Staresina et al., 2015; Klinzing et al., 2019), generalizes across brain networks.

### 4.2. SO–spindle coupling seen from the sources

The scalp and the sources differed in how spindle activity was organized around SOs. Slow-sigma activity peaked just before the SO trough at the scalp and in every region group, with closely matching timing. Fast-sigma activity did not: its rise into the SO up-state appeared only at the scalp, where it also began earlier than in any local source (Fig. 5), and the scalp–source difference had the same shape whichever region’s SOs were used. One explanation consistent with the coincidence results is that the scalp signal sums fast spindles from many regions, each loosely timed relative to a given region’s SOs, which would broaden and advance the averaged scalp rise. When slow and fast sigma were combined in a single 9–18 Hz band, their opposite preferences for the trough and the up-state cancelled, and no modulation remained at the scalp. This illustrates how a broad spindle band can obscure two distinct phase relationships, and supports analysing the two bands separately (or a finer-grained frequency analysis, as spindle frequency may be best characterized as a a gradient across the cortex, see Greenlaw et al. (2026)). More generally, the preferred phase or timing of spindle activity relative to SOs measured at the scalp should not be assumed to characterize the coupling of any individual brain region.

### 4.3. Amplitude and density are distinct properties

Amplitude and density maps were both highly reliable, yet shared little spatial variance, and the direction of their relationship depended on the oscillation: regions with larger fast spindles also had more of them, whereas SOs, and slow spindles once depth was accounted for, were larger where they were less frequent. Density and amplitude plausibly reflect different aspects of the underlying activity. Density may index how often the circuits that initiate events are engaged, whereas amplitude depends on the number and synchrony of contributing neurons, their spatial arrangement and orientation, and the extent to which an event spreads (Purcell et al., 2017; Cox et al., 2017; Knoblauch et al., 2003). Spectral measures, particularly from a single channel, combine the two, so that a change in sigma or delta power cannot be attributed to a change in how often events occur or in how large they are (Fig. 1). These differences were not explained by source depth: they persisted after partialling depth and within cortex, with the exception of the weak relationship for 2–4 Hz SOs.

### 4.4. Deep and cerebellar structures

SOs and spindles were detected in the thalamus, the hippocampus and amygdala, and the basal ganglia, at densities comparable to the cortical median. These observations are consistent with intracranial evidence of thalamic involvement in coordinating SOs and spindles (Schreiner et al., 2022), of spindle activity in the basal ganglia (Salih et al., 2009; Mizrahi-Kliger et al., 2022), and with MEG evidence of thalamocortical activity during spindles (Rattray et al., 2025). SO coincidence with the scalp was strongest in the hippocampus and amygdala and the thalamus. This should be interpreted cautiously given the small number of deep ROIs and the possibility of spatial leakage, but it suggests that scalp SOs are well aligned with SO activity in these structures.

The scalp–source difference in the onset of the fast-sigma rise was largest for the thalamus and the hippocampus and amygdala (about 0.42 s) and smallest for sensorimotor cortex (0.16 s; Supplementary Table S3). Because the scalp rise preceded the local rise in every group, this pattern is unlikely to reflect a fixed propagation delay from deep structures to the cortex; intracranial recordings suggest instead that thalamic spindles can lead cortical ones (Schreiner et al., 2022). It fits the composite account above, in which the further a region’s own spindle timing departs from the mixture recorded at Cz, the larger the apparent offset.

The cerebellum stood out in the spindle analyses. Fewer slow spindles were detected in the cerebellum than in the Cz–M1 EEG channel (event-rate ratio 0.85, against a median of 1.17 across ROIs), and cerebellar events coincided less with scalp events (coincidence strength 14.9% against a median of 17.8%). In the same dataset, spindle-band activity in the cerebellum was evident in spectral analyses (Greenlaw et al., 2026), and spindle-like activity has been recorded directly in the primate cerebellum (Xu et al., 2021). Event detection and spectral measures may capture different aspects of cerebellar activity, and whether the cerebellum generates its own spindles or follows thalamocortical ones remains open.

### 4.5. Implications for closed-loop stimulation

Closed-loop auditory stimulation can enhance or disrupt ongoing oscillations depending on timing (Ngo et al., 2013; Jourde et al., 2025), and it acts on whatever the detector sees. Knowing which circuits a scalp signature reflects therefore matters for both the intended and the unintended effects of stimulation (Fattinger et al., 2025). Our results show that a central derivation detects only a subset of regional SOs, and that the timing of SO-locked fast-sigma activity at the scalp does not match that of local sources. Stimulation timed to scalp events may therefore reach different circuits, at different phases, than assumed. Learning-based detectors such as the Portiloop already detect sleep events in real time from few channels (Valenchon et al., 2022). Building on such approaches, future detectors could be trained with MEG-derived labels to estimate the likely regional origin of an EEG-detected event, or to weight activity from particular regions, such as sensorimotor cortex, when targeting specific circuits.

### 4.6. Limitations

Several limitations should be considered. Data were collected during daytime naps, which mainly sample early-sleep dynamics and do not capture changes across a full night, such as the evolution of SO and spindle properties across sleep cycles. N2 and N3 epochs were pooled; stage-dependent regional differences in oscillatory power have been reported in this dataset (Greenlaw et al., 2026), and stage-specific maps of event amplitude and density remain to be established. Although the sample was modest, MEG studies of fundamental sleep physiology can be well powered with samples of this size (Baker et al., 2021), and our main inferences rest on participant-level tests. Source leakage and reduced sensitivity to deep sources are inherent to MEG source reconstruction, although recent work has distinguished activity from deep and cerebellar regions (Greenlaw et al., 2026; Rattray et al., 2025), and coincidence analyses are likely more exposed to leakage than the timing analyses. Detection thresholds were defined relative to each ROI and epoch, which makes detection robust to differences in signal strength between regions but yields higher event densities than fixed-threshold scalp detectors, and density should be interpreted as how often activity stands out from the local background. MEG polarity was aligned to Cz–M1 to resolve the sign ambiguity of source estimates; this fixes a common convention but does not establish whether polarity differences between regions are physiological or reconstruction-related. Finally, only a single scalp derivation was available for comparison. This configuration reflects common practice in polysomnography and closed-loop stimulation, in which one or a few EEG channels are used to characterize sleep neurophysiology or detect events for stimulation purposes; but other derivations, references and montages may capture different subsets of regional activity. SO–spindle coupling was examined only through SO-locked sigma activity, and other coupling measures were not explored.

### 4.7. Future directions

Building on these maps, future work could examine regional variation in SO–spindle coupling across different coupling definitions, and map how SOs propagate across regions, including their travelling patterns and delays. Studies combining full-cap EEG with MEG could test whether the relationships observed here for Cz–M1 extend to other derivations. MEG could also be used to train EEG-based detectors that identify oscillations by likely region of origin, allowing closed-loop stimulation to target events selectively and providing a causal test of their functional roles. The present recordings were part of a broader study that included a motor sequence learning task (Greenlaw et al., 2026), which offers an opportunity to relate regional SO and spindle activity to memory consolidation. The distinct cerebellar patterns observed in both this and the preceding work also warrant further investigation, as there is much to be learned about the cerebellum’s roles in sleep (Greenlaw et al., 2026; Xu et al., 2021; Canto et al., 2017).

### 4.8. Conclusion

Slow oscillations and sleep spindles occur throughout the brain, including deep and cerebellar structures, with amplitude and density forming distinct maps. A single scalp derivation samples a subset of local SOs and combines fast spindles from many regions into events whose timing does not match any one of them. How SOs and spindles are detected, and where, therefore shapes which circuits are studied and targeted. Whole-brain maps such as those presented here provide a reference for interpreting scalp findings and for designing more precise neuromodulation.

## Disclosure Statement

Financial disclosure: none. Non-financial disclosure: none.

## Data availability

Raw MEG and EEG data are available from the corresponding author on reasonable request, subject to approval by the research ethics boards.

## 5. Acknowledgements

EBJC was financially supported by grants from the FRQNT, Natural Sciences and Engineering Research Council of Canada (NSERC), and a Concordia University Research Chair in Sleep and Sound.

## CRediT authorship contribution statement

**Anne Calvel:** Formal analysis, Writing—original draft, Writing—review and editing, Visualization. **Keelin Greenlaw:** Data collection, Formal analysis. **Christopher J. Steele:** Methodology, Data collection, Writing-review and editing. **Emily B.J. Coffey:** Conceptualization, Methodology, Formal analysis, Writing-original draft, Writing-review and editing, Visualization, Supervision, Project administration.

## Notes

### Competing Interest Statement

The authors have declared no competing interest.

