## Supplementary figures (S1, S2, S3, S4, S5) and tables (S1, S2, S3) for "What EEG doesn’t see: a whole-brain MEG map of sleep spindles and slow oscillations"

### 1. Supplementary material

 (E.B.J. Coffey)
ORCID(s): 0000-0001-8249-7396 (E.B.J. Coffey)

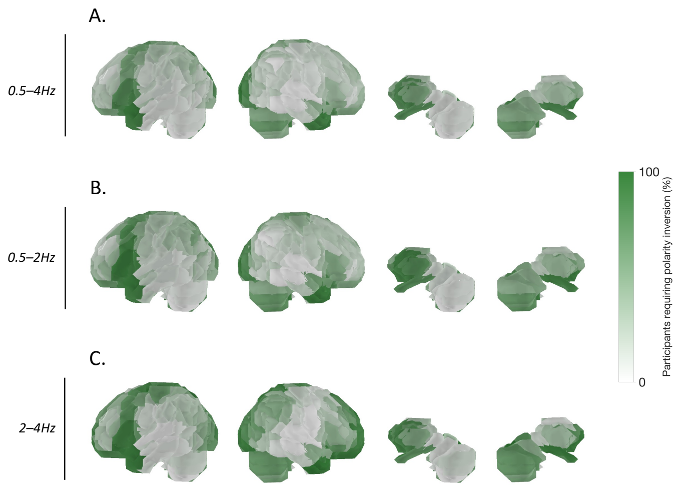


**Figure S1: Percentage of participants requiring MEG polarity inversion relative to EEG Cz.** (A) SOs detected in the 0.5–4 Hz band. (B) SOs detected in the 0.5–2 Hz band. (C) SOs detected in the 2–4 Hz band. For each ROI, color indicates the percentage of participants for whom the MEG and EEG Cz signals had opposite signs at the SO trough. White indicates 0% and dark green indicates 100%.

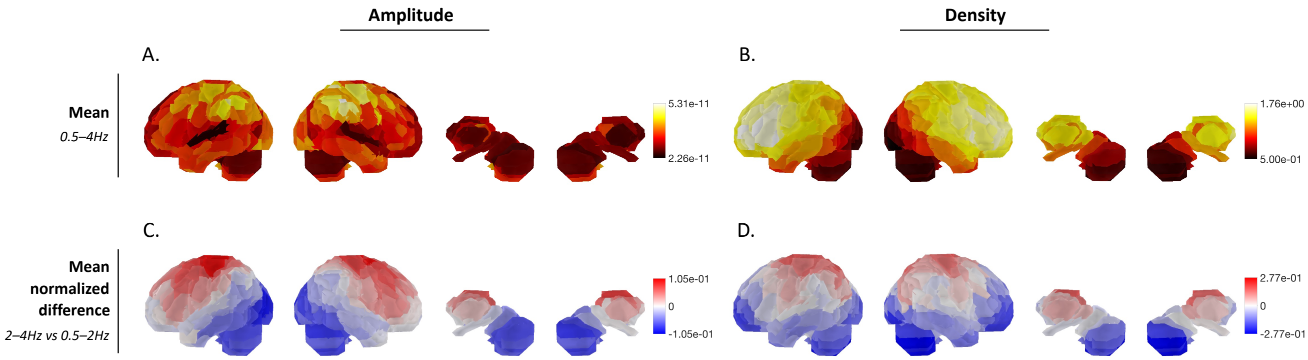


**Figure S2:** Whole-brain maps of spindles amplitude and density (group mean across participants, $N = 19$). A: 9–18 Hz spindles amplitude, defined as the mean peak RMS amplitude of detected spindles. B: 9–18 Hz density, defined as the number of detected spindles per minute of N2 and N3 sleep. For A–B, values were normalized within each map; colour bars show the original minimum and maximum values. C: Relative amplitude difference between the 13–18 Hz and 9–12.9 Hz bands. D: Relative density difference between the 13–18 Hz and 9–12.9 Hz bands. For C–D, relative differences were calculated as $(f - s)/(f + s)$, where $f$ and $s$ are the fast (13–18 Hz) and slow (9–12.9 Hz) values. Each map was centred on its brain-wide mean (red: 13–18 Hz relatively more prominent than the brain-wide average; blue: 9–12.9 Hz relatively more prominent). Brain-wide means: amplitude -0.11 (fast/slow ratio 0.81; regional range 0.66–1.00), density 0.00 (fast/slow ratio 1.0; regional range 0.6–1.5). Colour scales are set separately for each map.

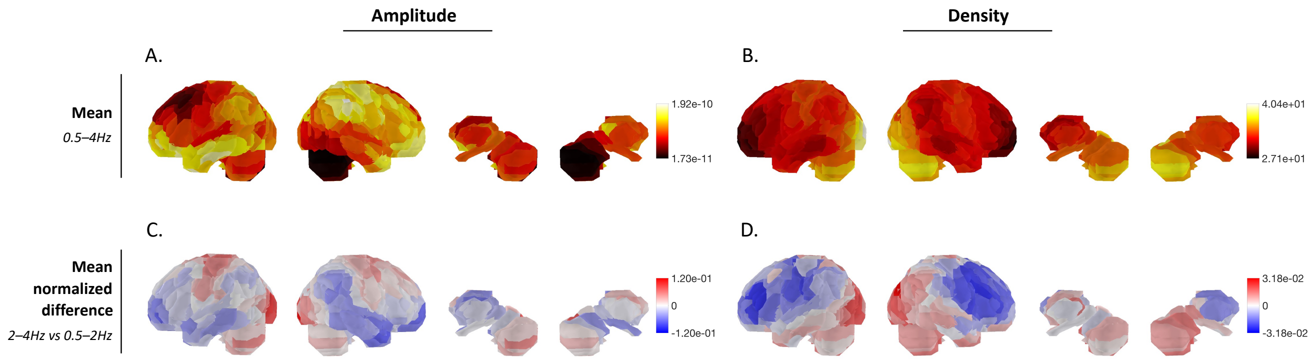


**Figure S3: Whole-brain maps of slow oscillation (SO) amplitude and density** (group mean across participants, $N = 19$). A: 0.5–4 Hz amplitude, defined as the mean trough-to-peak amplitude of detected SOs. B: 0.5–4 Hz density, defined as the number of detected SOs per minute of N2 and N3 sleep. For A–B, values were normalized within each map; colour bars show the original minimum and maximum values. C: Relative amplitude difference between the 2–4 Hz and 0.5–2 Hz bands. D: Relative density difference between the 2–4 Hz and 0.5–2 Hz bands. For C–D, relative differences were calculated as $(f - s)/(f + s)$, where $f$ and $s$ are the 2–4 Hz and 0.5–2 Hz values. Each map was centred on its brain-wide mean (red: 2–4 Hz relatively more prominent than the brain-wide average; blue: 0.5–2 Hz relatively more prominent). Brain-wide means: amplitude -0.33 (2–4 Hz SOs about half the amplitude), density 0.48 (2–4 Hz SOs about 2.8 times as frequent). Colour scales are set separately for each map; the regional variation in the density ratio is small (fast/slow ratio 2.7–3.1) compared with amplitude (0.43–0.66).

**Table S1**

Regions of interest included in each anatomical group. Each listed region was represented bilaterally. Numbers in parentheses indicate the included atlas subdivisions.

| Region group | $n$ | Included ROIs |
|---|---|---|
| Sensorimotor cortex | 34 | Paracentral lobule (1–4), postcentral sulcus (1–3), precentral sulcus (1–6), and central sulcus/Rolando (1–4). |
| Occipital cortex | 50 | Calcarine cortex (1–3), cuneus (1–2), lingual gyrus (1–6), inferior occipital gyrus (1–2), lateral occipital gyrus (1–5), middle occipital gyrus (1–4), occipital pole (1), and superior occipital gyrus (1–2). |
| Remaining cortex | 256 | Angular gyrus (1–3); anterior, middle, and posterior cingulate gyri (1–2, 1–3, and 1–3); inferior orbital frontal gyrus (1–2); inferior triangular frontal gyrus (1); medial orbital frontal gyrus (1–2); middle frontal gyrus (1–5); middle orbital frontal gyrus (1–2); superior frontal gyrus (1–3); superior medial frontal gyrus (1–3); superior orbital frontal gyrus (1); fusiform gyrus (1–7); anterior insula (1–5); posterior insula (1); parahippocampal gyrus (1–5); inferior parietal gyrus (1); superior parietal gyrus (1–5); precuneus (1–9); Rolandic operculum (1–2); subcallosal gyrus (1); supplementary motor area (1–3); supramarginal gyrus (1–7); inferior temporal gyrus (1–5); middle temporal gyrus (1–4); middle temporal pole (1–3); superior temporal pole (1–2); superior temporal gyrus (1–4); anterior rostral sulcus (1); cingulate sulcus (1–7); inferior frontal sulcus (1–2); intraoccipital sulcus (1); intraparietal sulcus (1–3); olfactory sulcus (1); orbital sulcus (1–2); parieto-occipital sulcus (1–6); superior frontal sulcus (1–6); and superior temporal sulcus (1–5). |
| Thalamus | 18 | Thalamus (1–9). |
| Hippocampus and amygdala | 6 | Hippocampus (1–2) and amygdala (1). |
| Other subcortical nuclei | 20 | Caudate nucleus (1–7), pallidum (1), and putamen (2–3). |
| Cerebellum | 32 | Cerebellar lobules 3, 4–5, 6, 7b, 8, 9, and 10; Crus I and Crus II; and vermian lobules 1–2, 3, 4–5, 6, 7, 8, 9, and 10. Vermian lobules 3 and 8 were represented by single midline ROIs; all other entries were bilateral. |

**Table S2**

Amplitude–density correspondence: control analyses for source geometry. Values are attenuation-corrected Spearman $\rho$ [95% CI].

| Band | $\rho$(amp., depth) | $\rho$(dens., depth) | All ROIs (416) | Depth partialled (416) | Cortex only (340) |
|---|---|---|---|---|---|
| SO 0.5–2 Hz | −0.24 | 0.24 | −0.41 [−0.51, −0.22] | −0.36 [−0.47, −0.17] | −0.34 [−0.46, −0.12] |
| SO 2–4 Hz | −0.28 | 0.27 | −0.15 [−0.24, −0.05] | −0.08 [−0.20, 0.01] | −0.05 [−0.15, 0.06] |
| SO 0.5–4 Hz | −0.25 | 0.23 | −0.43 [−0.51, −0.24] | −0.39 [−0.48, −0.22] | −0.36 [−0.47, −0.16] |
| Spindles, slow | −0.46 | −0.29 | −0.22 [−0.39, 0.08] | −0.44 [−0.60, −0.08] | −0.26 [−0.45, 0.09] |
| Spindles, fast | −0.49 | 0.06 | 0.35 [0.11, 0.48] | 0.44 [0.16, 0.60] | 0.37 [0.12, 0.51] |
| Spindles, all | −0.52 | −0.19 | −0.04 [−0.25, 0.23] | −0.18 [−0.37, 0.16] | −0.08 [−0.29, 0.23] |

Depth: distance from the ROI centroid (MNI) to the outer surface of the template brain; depth correlations are across all 416 ROIs. Depth partialled: all correlations, reliabilities and the participant-level test computed as partial Spearman correlations controlling for depth. Participant-level same- vs cross-measure tests: all $t(18) \geq 15.7$, Holm-corrected $p < 10^{-11}$ in each analysis.

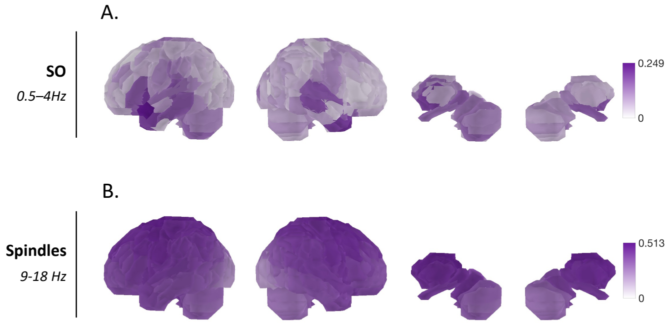


**Figure S4: Spatial distribution of continuous EEG–MEG correlations.** (A) Correlations between continuous EEG Cz–M1 and polarity-corrected MEG signals in each ROI in the SO band (0.5–4 Hz). (B) Correlations between continuous RMS amplitude envelopes of EEG Cz–M1 and MEG signals in each ROI in the spindle band (9–18 Hz). Pearson correlations were calculated for each 30-s N2/N3 epoch, Fisher-$z$ transformed, averaged across epochs within each participant and then across participants, and back-transformed to $r$ for visualization. White indicates $r = 0$, and darker purple indicates stronger positive correlations.

**Table S3**

Jackknife latency estimates of SO-locked sigma activity in the local MEG source and at Cz–M1, for SOs (0.5–2 Hz) detected in each region group ($N = 19$). Times are in seconds relative to the local SO trough.

| | Fast sigma: rise onset (half-maximum) | | | | Slow sigma: peak | | | | SO trough |
|---|---|---|---|---|---|---|---|---|---|
| Region group | MEG | EEG | EEG−MEG | $p$ | MEG | EEG | EEG−MEG | $p$ | EEG−MEG (ms) |
| Sensorimotor | −0.49 | −0.65 | −0.16 | 0.025 | −0.053 | −0.080 | −0.027 | 0.072 | 0 |
| Occipital | −0.47 | −0.81 | −0.34 | <0.001 | −0.087 | −0.080 | 0.007 | 0.842 | 0 |
| Remaining cortex | −0.45 | −0.78 | −0.33 | <0.001 | −0.073 | −0.100 | −0.027 | 0.028 | −13 |
| Thalamus | −0.43 | −0.85 | −0.42 | <0.001 | −0.073 | −0.120 | −0.047 | 0.311 | −20 |
| Hippocampus & amygdala | −0.42 | −0.84 | −0.42 | <0.001 | −0.073 | −0.153 | −0.080 | 0.192 | −27 |
| Other subcortical | −0.43 | −0.74 | −0.31 | 0.047 | −0.093 | −0.100 | −0.007 | 0.697 | −20 |
| Cerebellum | −0.41 | −0.60 | −0.19 | 0.278 | −0.133 | −0.120 | 0.013 | 0.838 | −20 |

Latencies were measured on the group-average envelopes. Standard errors of the EEG–MEG differences were estimated from leave-one-participant-out averages (jackknife; see Methods), and differences were tested as $t$ = difference / jackknife SE (df = 18; uncorrected $p$). Fast-sigma rise onset: time at which the envelope reached half of its rise from the minimum in −1.0 to −0.4 s to the maximum in −0.4 to 0.1 s. Slow-sigma peak: time of the maximum in −0.5 to 0.2 s. SO trough: lag of the Cz–M1 SO trough relative to the local MEG trough at the same events; negative values indicate that the EEG trough came first.

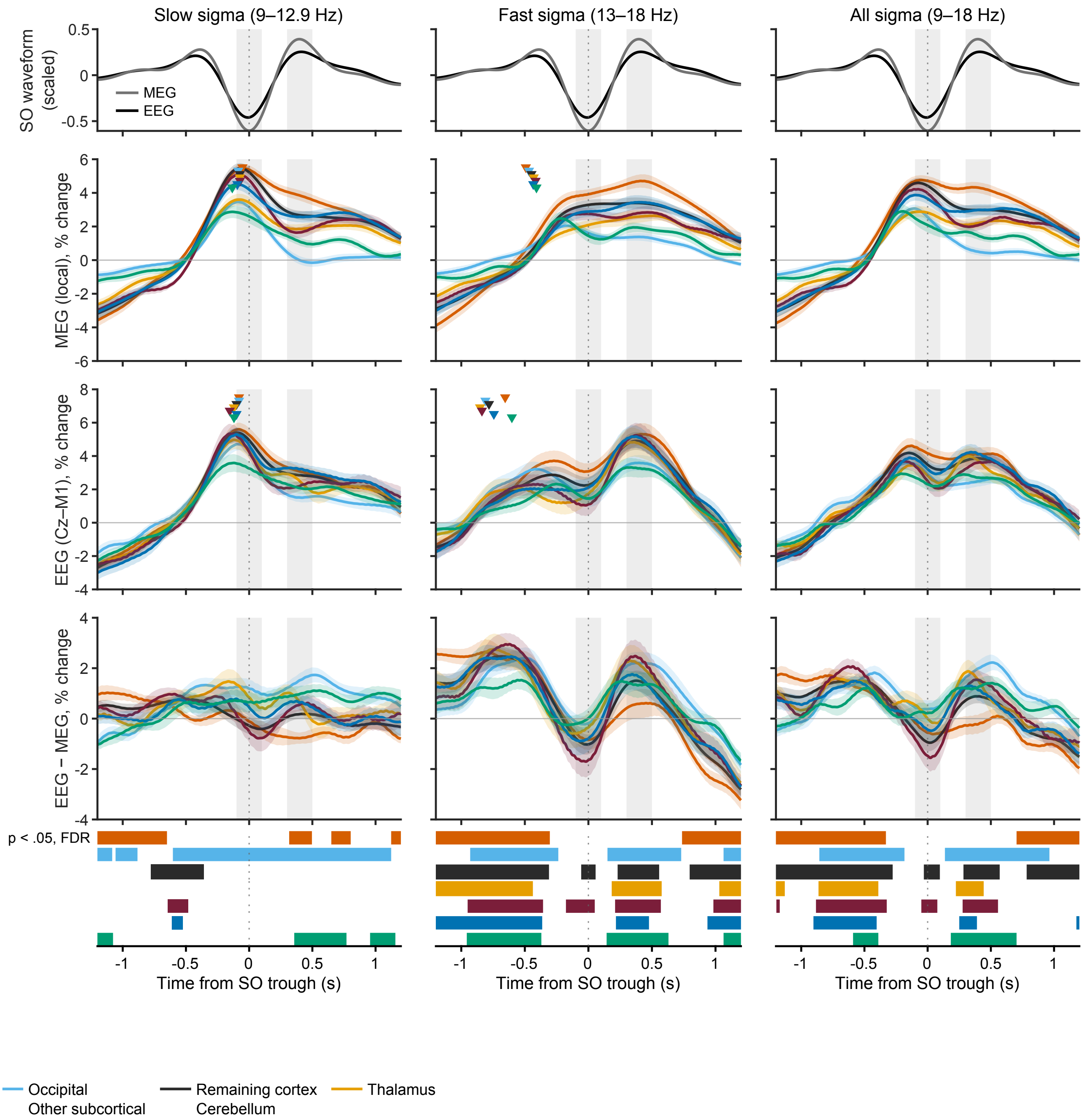


**Figure S5:** SO-locked sigma activity for all seven region groups (SOs detected in each region, 0.5–2 Hz). Columns: slow (9–12.9 Hz), fast (13–18 Hz) and all (9–18 Hz) sigma. Top row: SO waveform averaged over all ROIs (MEG, grey; EEG, black; each scaled to its range). Second row: local MEG source activity; third row: EEG (Cz–M1) at the same events; fourth row: EEG minus MEG. Sigma is expressed as percent change from its mean over $\pm 2$ s and averaged over the ROIs of each group: sensorimotor cortex (vermillion), occipital cortex (light blue), remaining cortex (dark grey), thalamus (orange), hippocampus and amygdala (burgundy), other subcortical nuclei (dark blue) and cerebellum (green). Shading, $\pm 1$ SEM across participants ($N = 19$). Grey bands mark the trough and up-state windows derived from the group-average SO waveform. Triangles in the MEG and EEG rows mark jackknife latency estimates for each group (slow sigma: peak; fast sigma: half-maximum of the pre-trough rise). Strips below the bottom row mark, for each group, the time points at which EEG and MEG differ (paired $t$-test at each sample, FDR-corrected across samples within $\pm 1.2$ s). Group membership is listed in Supplementary Table S1, and latency estimates in Supplementary Table S3.
